# Analysis of socioeconomic disadvantage and pace of aging measured in saliva DNA methylation of children and adolescents

**DOI:** 10.1101/2020.06.04.134502

**Authors:** Laurel Raffington, Daniel W. Belsky, Margherita Malanchini, Elliot M. Tucker-Drob, K. Paige Harden

## Abstract

Children who grow up in socioeconomically disadvantaged families face increased burden of disease and disability as they mature into adulthood. One hypothesized mechanism for this increased burden is that early-life disadvantage and its associated psychological stress accelerate biological processes of aging, increasing vulnerability to subsequent disease. In order to evaluate this hypothesis and the potential impact of preventive interventions, measures to quantify the early acceleration of biological aging in childhood are needed. Here, we evaluated a novel DNA-methylation measure of the pace of aging, DunedinPoAm, and compared DunedinPoAm results with results for several published epigenetic clocks. Data on saliva DNA-methylation and socioeconomic circumstances were collected from *N* = 600 children and adolescents aged 8- to 18-years-old (48% female) participating in the Texas Twin Project. Participants living in more disadvantaged families and neighborhoods exhibited faster pace of aging (*r* = 0.18, *p* = 0.001 for both). Latinx-identifying children exhibited faster DunedinPoAm compared to both White- and Latinx-White-identifying children, consistent with higher levels of disadvantage in this group. Children with more advanced pubertal development and those with had higher body-mass index also exhibited faster DunedinPoAm, but these covariates did not account for the observed socioeconomic gradient in methylation pace of aging. In contrast to findings for DunedinPoAm, we did not detect associations of socioeconomic disadvantage with five published epigenetic clocks. Findings suggest that DNA-methylation pace-of-aging measures may prove more sensitive to health damaging effects of adversity, particularly when measurements are taken early in the life course, before substantial aging has occurred.

---

Individuals who are exposed to social adversity in childhood experience a higher burden of aging-related disease later in life (1). In particularly, children raised in conditions of low socioeconomic status, and who thus experience a suite of material hardships and psychological stressors, are at increased risk for wide range of later-life health problems, including cardiovascular disease, Type-II diabetes, cancer, anxiety, and dementia, as well as shorter lifespan (2–4). These childhood socioeconomic gradients in adult-onset disorders partly reflect socioeconomic gradients in health problems that onset during childhood, including obesity, asthma, and stress-related mental health problems (5–7). However, adult health continues to be graded by childhood socioeconomic status, even after accounting for childhood-onset health problems and for adult socioeconomic status (2,6,8). This social patterning is also observed in other animals, including species in whom childhood social conditions and adult social conditions are less strongly correlated than in humans (1).

A leading hypothesis for the enduring childhood socioeconomic gradient in adult health is that social disadvantage initiates biological changes in children that ultimately make them more vulnerable to developing disease in adulthood (9,10). The link from childhood disadvantage to multiple different disease processes in adulthood suggests the further hypothesis that these biological changes involves an acceleration of the process of biological aging (11,12). In adults, the social determinants of health and aging have been studied using multi-organ system measures such as allostatic load (13) and, more recently, physiology-based measures of biological aging (14). Adults who grew up under conditions of socioeconomic disadvantage exhibit a faster pace of aging and more advanced biological age according to these measures (11,15).

Yet, measures taken in adulthood are decades removed from the early exposures thought to be critical for shaping health gradients. Methods to quantify processes of biological aging in children are needed. A barrier to implementing physiology-based indices in studies of childhood is that processes of growth and development during childhood may confound physiological measurements originally designed to capture stress- and aging-related decline in organ system integrity in adults. Moreover, intensive and/or invasive multimodal measures are difficult to collect from large samples of children. Methods that can quantify processes of biological aging from a single accessible tissue, such as saliva, are a priority for research with child populations.

Molecular measures that aim to quantify the aging process at the cellular level provide an alternative approach for investigating the developmental roots of adult health risk. Telomere length, a biomarker of cellular aging, is one candidate (16,17). But questions remain about whether telomere length measured in saliva or blood constitutes a valid biomarker of aging at the organism level (18,19). More recently, DNA-methylation measures, called “epigenetic clocks,” have emerged as leading measures of biological aging in humans and other species (20). Epigenetic clocks are algorithms that estimate a person’s age from dozens or hundreds of DNA-methylation marks across the genome. Clock-estimated ages are highly correlated with chronological ages (*e.g.*, Pearson *r* > 0.9 in mixed-age samples). The difference between a person’s clock-estimated age and their true chronological age, *aka* “age acceleration,” is proposed as a measure of biological aging (20).

In adults, some epigenetic clocks detect evidence of more advanced aging associated with socioeconomic disadvantage (21). In children, however, findings are mixed (12,22). Not all types of adversity show evidence of association with clock measurements. In particular, low childhood socioeconomic status, which is consistently associated with shorter healthy lifespan, is not consistently associated with epigenetic clock measures of aging (23).

An alternative to epigenetic clocks is a new DNA methylation-based measurement of the pace of biological aging, DunedinPoAm (24). Whereas the clocks were developed from analysis comparing chronologically older individuals to younger ones, DunedinPoAm was developed from analysis of change over time occurring in a cohort of individuals who were all the same chronological age. In contrast to epigenetic clocks, which aim to quantify the *amount* of aging an individual has experienced up to the time of measurement, *pace* of aging measures aim to quantify how fast the individual is aging (25). Because children have not lived very long, they may not yet have accumulated large differences in how much aging has occurred by the time of measurement. Epigenetic clocks may, therefore, be less sensitive to changes in biological aging as compared to pace of aging measures such as DunedinPoAm. One previous study showed that DunedinPoAm indicates faster methylation pace of aging in 18 year-olds exposed to early adversity, including low childhood socioeconomic status (24). However, this new measure has not yet been studied in children.

We analyzed saliva DNA-methylation from 600 White- or Latinx-identifying children aged 8-18 from the population-based Texas Twin Project to examine whether family-level and neighborhood-level cumulative socioeconomic disadvantage were associated with faster methylation pace of aging. We also examined whether children’s racial/ethnic identities were associated with methylation pace of aging. For comparison, we repeated analysis using several epigenetic clocks **(26–30)**. We conducted additional analysis to evaluate how smoking, body mass index, and pubertal status may affect associations of childhood socioeconomic disadvantage with proposed DNA-methylation measures of biological aging.

## Methods and Materials

### Participants

Participants were 600 (285 female) children and adolescents from 328 unique families aged 8 to 18 years (*M* = 12.68, *SD* = 3.02) from the Texas Twin Project (31). The Texas Twin Project is an ongoing longitudinal study that includes the collection of salivary samples. Saliva samples were selected to be assayed for DNA methylation using EPIC arrays if participants self-identified their race/ethnicity as White and/or Latinx and had contributed cortisol data (not reported here). After excluding 8 participants during DNA-methylation preprocessing, there were *N* = 457 participants who identified as only White, *N* = 77 as only Latinx, and *N* = 61 as Latinx and White. We capitalize these terms to highlight that racial/ethnic identities are social constructions that are not based on “innate” biosocial boundaries, but may have biosocial effects through people’s lived experiences (32).

### DNA-methylation

Saliva samples were collected during a laboratory visit using Oragene kits (DNA Genotek, Ottawa, ON, Canada). DNA extraction and methylation profiling were conducted by Edinburgh Clinical Research Facility (UK). The Infinium MethylationEPIC BeadChip kit (Illumina, Inc., San Diego, CA) was used to assess methylation levels at 850,000 methylation sites. Methylation profiles were residualized for cell composition, array and slide; all samples came from the same batch. See Supplement for DNA methylation preprocessing details and cell composition estimation and correction.

### DunedinPoAm

DunedinPoAm was calculated based on the published algorithm (24) using code available at https://github.com/danbelsky/DunedinPoAm38. Briefly, DunedinPoAm was developed from DNA-methylation analysis of Pace of Aging in the Dunedin Study birth cohort. Pace of Aging is a composite phenotype derived from analysis of longitudinal change in 18 biomarkers of organ-system integrity measured when Dunedin Study members were all 26, 32, and 38 years of age (25). Elastic-net regression machine learning analysis was used to fit Pace of Aging to Illumina 450k DNA-methylation data generated from blood samples collected when participants were aged 38 years. The elastic net regression produced a 46-CpG algorithm. Increments of DunedinPoAm correspond to “years” of physiological change occurring per 12-months of chronological time. The Dunedin Study mean was 1, *i.e.* the typical pace of aging among 38-year-olds in that birth cohort. Thus, 0.01 increment of DunedinPoAm corresponds to a percentage point increase or decrease in an individual’s pace of aging relative to the Dunedin birth cohort at midlife.

### Epigenetic clocks

We computed five epigenetic clocks. The original clocks proposed by Horvath and by Hannum et al. were derived from DNA-methylation analysis of chronological age (26,27). The same approach was used to develop a pediatric clock optimized to predict the age of children from buccal cell DNA methylation (PedBE) (30). In addition to these three chronological-age-based clocks, we analyzed two recently published clocks developed from DNA-methylation analysis of mortality risk, PhenoAge and GrimAge (28,29). These clocks were developed in two steps. In a first step, mortality risk was modeled from chronological age and a panel of biomarkers. In the second step, predicted risk derived from the first-step model was in turn modeled from DNA-methylation data. These clocks remain highly correlated with chronological age, but are more strongly related to disease and mortality as compared to the Horvath and Hannum et al. clocks (29). The Horvath clock was developed from analysis of multiple tissues. The Hannum clock and the PhenoAge and GrimAge clocks were developed from analysis of blood DNA methylation. The PedBE clock was developed from analysis of buccal cells.

Epigenetic clocks were computed using the web-based tool hosted by the Horvath Lab (https://dnamage.genetics.ucla.edu/home). Following standard methods, we converted clocks to age-acceleration residuals for analysis by regressing participants’ computed epigenetic-clock age values on their chronological ages and predicting residual values.

### Cumulative socioeconomic disadvantage

We measured children’s socioeconomic disadvantage at the family and neighborhood levels of analysis.

#### Family-level socioeconomic disadvantage

The family-level measure was computed from parent reports of household income, parental education, occupation, history of financial problems, food insecurity (based on the US Household Food Security Survey Module, 31), father absence, residential instability (changes in home address), and family receipt of public assistance. These were aggregated to form a composite measure of household-level cumulative socioeconomic disadvantage (*M* = -0.08, *SD* = 0.89), which is slightly below the larger sample’s mean described in (34), and coded such that higher scores reflect greater disadvantage.

#### Neighborhood-level socioeconomic disadvantage

The neighborhood-level measure was composed from tract-level US Census data according to the method described in (34). Briefly, participant addresses were linked to tract-level data from the US Census Bureau American Community Survey averaged over five years (https://www.census.gov/programs-surveys/acs). A composite score of neighborhood-level socioeconomic disadvantage was computed from tract-level proportions of residents reported as unemployed, living below the Federal poverty threshold, having less than 12 years of education, not being employed in a management position, and single mothers. The average neighborhood-level socioeconomic disadvantage (*M* = -0.09, *SD* = 0.87) was slightly below the larger sample’s mean described in (34), and coded such that higher scores reflect greater disadvantage.

### Health behavior covariates

Smoking and obesity are socially-patterned health behavior exposures that are more common in children from lower socioeconomic status families and neighborhoods (7,35). They are also associated with differential DNA-methylation patterns across the genome (36–38). We therefore considered tobacco exposure and body-mass index in our analysis.

#### Tobacco exposure

We measured tobacco exposure from (1) participant self-report of tobacco use, (2) a whole-genome DNA-methylation (poly-DNAm) smoking score (*M* = 0, *SD* = 0.33; (39)), and (3) methylation of the AHRR gene (*M* = 0, *SD* = 0.03; cg05575921 (40)).

#### Body mass index (BMI)

We measured BMI from in-laboratory measurements of height and weight transformed to sex- and age-normed z-scores according to the method published by the US Centers for Disease Control and Prevention (*M* = 0.3, *SD* = 1.32; https://www.cdc.gov/growthcharts/percentile_data_files.htm).

### Pubertal development

Age-adjusted pubertal status is proposed as an index of biological aging in children (41). Puberty tends to onset at younger ages in children growing up in conditions of socioeconomic disadvantage (42,43). Puberty is also associated with a range of DNA-methylation changes (44,45). We therefore considered children’s pubertal development in our analysis.

#### Pubertal development

We measured pubertal development using children’s self-reports on the Pubertal Development Scale (46). The scale assesses the extent of development across five sex-specific domains (for both: height, body hair growth, skin changes; for girls: onset of menses, breast development; for boys: growth in body hair, deepening of voice). A total pubertal status score was computed as the average response (1 = “Not yet begun” to 4 = “Has finished changing”) across all items (*M* = 2.39, *SD* = 0.93). Pubertal development was residualized for age, sex, and an age by sex interaction. We also examined menarcheal status in girls (menses *N* = 153 girls; no menses *N* = 134 girls).

### Analysis

We tested associations of childhood socioeconomic disadvantage and racial/ethnic identity with the pace of biological aging using regression analysis. To account for non-independence of data on siblings, we fitted regressions using linear mixed models implemented with the lme4 *R* package. We report parameter estimates with bootstrapped 95% confidence intervals (computed with 500 simulations using lme4’s “confint.merMod”). Continuous measures were standardized for analysis to *M* = 0, *SD* = 1 allowing for interpretation of effect-sizes in the metrics of Pearson’s *r* in the case of continuously distributed exposure variables and Cohen’s *d* in the case of nominal variables (*i.e.*, race/ethnicity). All models were adjusted for sex and children’s chronological age.

To test if associations between socioeconomic disadvantage and pace of biological aging were accounted for by social gradients in tobacco exposure and/or obesity, we conducted covariate-adjusted regressions to evaluate sensitivity of results. We conducted parallel analysis to evaluate independence of associations from pubertal development.

For comparison, we repeated pace of aging analysis using age-acceleration residuals from five published epigenetic clocks.

## Results

DunedinPoAm measured from salivary DNA was approximately normally distributed in Texas Twin Project children and adolescents (before correction for the cell composition of saliva samples, mean DunedinPoAm was 0.79, *SD* = 0.05). DunedinPoAm was not correlated with age in this young sample (*r* = 0.015, *SE* = 0.047, 95% CI = -0.079 – 0.108, *p* = 0.747) and was similar in boys and girls (*d* = -0.029, *SE* = 0.0432, 95% CI = -0.107 – 0.059, *p* = 0.482).

### Socioeconomic disadvantage and Latinx racial/ethnic identity are associated with faster methylation pace of aging

We first tested if children growing up in more socioeconomically disadvantaged circumstances exhibited faster methylation pace of aging. We conducted separate analysis of socioeconomic disadvantage measured for children’s families and their neighborhoods. At both levels of analysis, children growing up under conditions of greater socioeconomic disadvantage exhibited faster pace of aging as measured by DunedinPoAm (family-level *r* = 0.176, *SE* = 0.050, 95% CI = 0.080 – 0.270, *p* = 0.001; neighborhood level *r* = 0.176, *SE* = 0.053, 95% CI = 0.073 – 0.277, *p* = 0.001; Figure 1).

**Figure 1.**
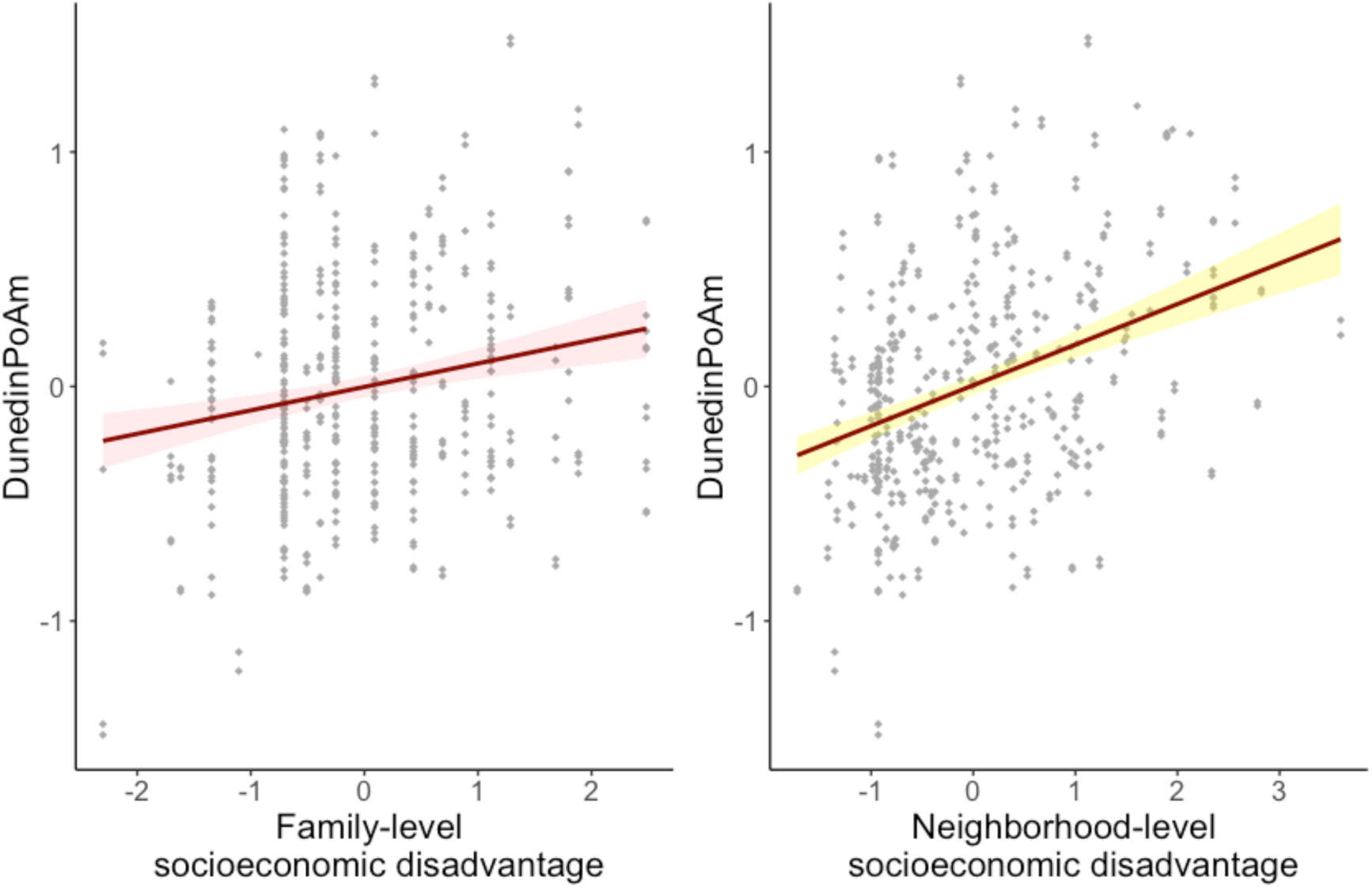
Associations between family-level and neighborhood-level socioeconomic disadvantage and DunedinPoAm. DunedinPoAm and socioeconomic disadvantage values are in standard deviation units. Higher values indicate a methylation profile of faster biological aging. Regression is estimated from linear mixed effects model that accounts for nesting of children within families. The shaded areas represent the smoothed lower and upper 95% confidence intervals.

We next tested if Latinx (12.9% of sample) and Latinx-White (10.3% of sample) identifying children exhibited faster DunedinPoAm, as compared to children identifying solely as White (76.8% of sample). Latinx-identifying children exhibited faster DunedinPoAm compared to both White-identifying children (*d* = -0.206, *SE* = 0.059, 95% CI = -0.319 – -0.092, *p* = 0.001) and Latinx-White identifying children (*d* = -0.159, *SE* = 0.059, 95% CI = -0.269 – -0.043, *p* = 0.008; Figure 2).

**Figure 2.**
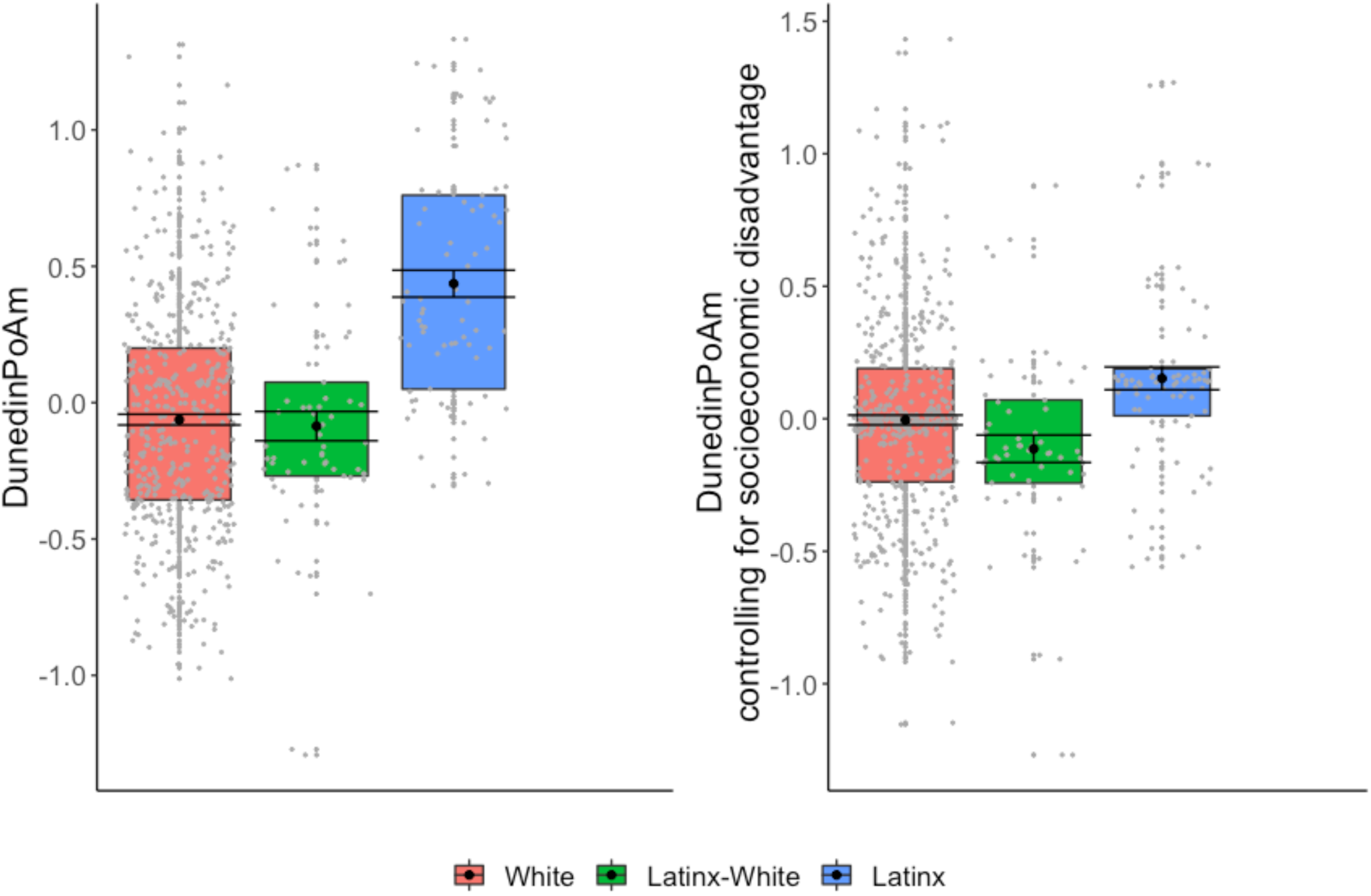
DunedinPoAm in children identifying as White, Latinx, and Latinx-White. DunedinPoAm values are in standard deviation units. Higher values indicate a methylation profile of faster biological aging. Regression is estimated from linear mixed effects model that accounts for nesting of children within families. The boxplot displays group DunedinPoAm differences in the mean (black circle), standard errors of the mean (error bars), and the first and third quartiles (lower and upper hinges). Group differences were significant at the alpha=0.05 threshold without adjustment for differences in socioeconomic disadvantage between groups (left panel), but were no longer significantly different from zero when controlling for family-level and neighborhood-level disadvantage (right panel).

Latinx children tended to be exposed to higher rates of family-level and neighborhood-level cumulative socioeconomic disadvantage compared to both White-identifying children (family-level: *d* = -0.329, *SE* = 0.074, 95% CI = -0.473. – -0.184, *p* < 0.001; neighborhood-level: *d* = -0.497, *SE* = 0.076, 95% CI = -0.548 – -0.295, *p* < 0.001) and Latinx-White identifying children (family-level: d = -0.208, *SE* = 0.072, 95% CI = -0.349 – -0.066, *p* = 0.004; neighborhood-level: d = -0.292, *SE* = 0.075, 95% CI = -0.350 – -0.111, *p* < 0.001). We therefore tested if racial/ethnic group differences in DunedinPoAm were statistically explained by socioeconomic differences between the groups of children. Adjusting for group differences in family-level and neighborhood-level disadvantage largely accounted for differences in methylation pace of aging between Latinx-identifying children and White-only identifying children (*d* = -0.068, *SE* = 0.081, 95% CI = -0.224 – 0.080 *p* = 0.406) or Latinx-White identifying children (*d* = -0.094, *SE* = 0.078, 95% CI = -0.225 – 0.046, *p* = 0.229; Figure 2); associations were no longer statistically different from zero. Statistical adjustment for racial/ethnic identity only modestly attenuated associations of socioeconomic disadvantage with DunedinPoAm; associations remained statistically different from zero (family-level: *r* = 0.158, *SE* = 0.053, 95% CI = 0.046 – 0.251, *p* = 0.003; neighborhood-level: *r* = 0.139, *SE* = 0.057, 95% CI = 0.033 – 0.256, *p* = 0.015).

### Associations between DunedinPoAm and socioeconomic disadvantage were robust to health behavior and developmental covariates

#### Smoking

To test if associations between socioeconomic disadvantage and pace of aging could be explained by differences in tobacco exposure between low and high disadvantage children, we conducted two sets of analyses. First, we repeated regression analysis excluding self-reported smokers. There were few self-reported smokers in the sample (*N* = 11 children, 1.8% of the sample). Results were unchanged excluding these participants. Second, we repeated regression analysis adding covariate adjustment for DNA-methylation measures of tobacco exposure (genome-wide smoking methylation (poly-DNAm) and AHRR smoking scores). Poly-DNAm smoking profiles were positively associated with DunedinPoAm (*r* = 0.194, SE = 0.042, 95% CI = 0.112 – 0.276, *p* < 0.001). Covariate adjustment for poly-DNAm and AHRR measures of tobacco exposure only modestly attenuated associations at the family-level (*r* = 0.159, *SE* = 0.050, 95% CI = 0.061 – 0.249, *p* = 0.001) and the neighborhood-level (*r* = 0.154, *SE* = 0.052, 95% CI = 0.053 – 0.254, *p* = 0.003).

#### BMI

To test if associations between socioeconomic disadvantage and pace of aging could be explained by differences in obesity/overweight, we repeated regression analysis adding covariate adjustment for BMI. Children with higher BMI had faster DunedinPoAm (*r* = 0.264, *SE* = 0.042, 95% CI = 0.178 – 0.345, *p* < 0.001). Covariate adjustment for BMI modestly attenuated associations of DunedinPoAm with socioeconomic disadvantage (family-level *r* = 0.138, *SE* = 0.049, 95% CI = 0.042 – 0.227, *p* = 0.005; neighborhood-level *r* = 0.115, *SE* = 0.052, 95% CI = 0.016 – 0.247, *p* = 0.027).

#### Pubertal development

To test if associations between socioeconomic disadvantage and methylation pace of aging could be explained by accelerated pubertal development in more disadvantaged children, we repeated regression analysis adding covariate adjustment for pubertal development. We considered two measures of puberty, the Pubertal Development Status scale and, in girls, menarcheal status. DunedinPoAm was weakly associated with self-reported pubertal development according to the Pubertal Development Scale, but the effect-size was not statistically different from zero (*r* = 0.070, *SE* = 0.040, 95% CI = -0.016 – 0.147, *p* = 0.083). DunedinPoAm indicated somewhat faster aging in girls who had experienced first menses as compared to those who had not. The effect-size was small, but statistically different from zero (*d* = 0.186, *SE* = 0.092, 95% CI = 0.004 – 0.379, *p* = 0.045).

Covariate adjustment for the pubertal development scale modestly attenuated associations of socioeconomic disadvantage with DunedinPoAm (family-level *r* = 0.173, *SE* = 0.051, 95% CI = 0.071 – 0.267, *p* = 0.001; neighborhood-level *r* = 0.158, *SE* = 0.054, 95% CI = 0.051 – 0.267, *p* = 0.003). Results were similar for covariate adjustment for menarcheal status among girls (unadjusted family-level *r* = 0.131, *SE* = 0.070, 95% CI = -0.004 – 0.263, *p* = 0.063; unadjusted neighborhood-level *r* = 0.182, *SE* = 0.070, 95% CI = 0.056 – 0.337, *p* = 0.010; adjusted family-level r = 0.135, *SE* = 0.073, 95% CI = -0.007 – 0.267, *p* = 0.066; adjusted neighborhood-level *r* = 0.171, *SE* = 0.072, 95% CI = 0.036 – 0.329, *p* = 0.020).

### Comparison of results with epigenetic clocks

We compared results for DunedinPoAm with results from analysis of five published epigenetic clocks: (1) Horvath DNAm age (before correction for the cell composition of saliva samples *M* = 14.96, *SD* = 4.38, 95% CI = 14.61-15.31), (2) Hannum DNAm age (before cell correction *M* = 22.57, *SD* = 3.46, 95% CI = 22.29 – 22.84), (3) PedBE (before cell correction *M* = 11.04, *SD* = 1.98, 95% CI = 10.88 – 11.19), (4) PhenoAge (before cell correction *M* = 17.26, *SD* = 5.53, 95% CI = 16.82 – 17.70), and (5) GrimAge (before cell correction *M* = 35.84, *SD* = 3.33, 95% CI = 35.58 – 36.11). All five epigenetic clocks were strongly correlated with chronological age (median *r* = 0.74, Figure 3, panel A).

**Figure 3.**
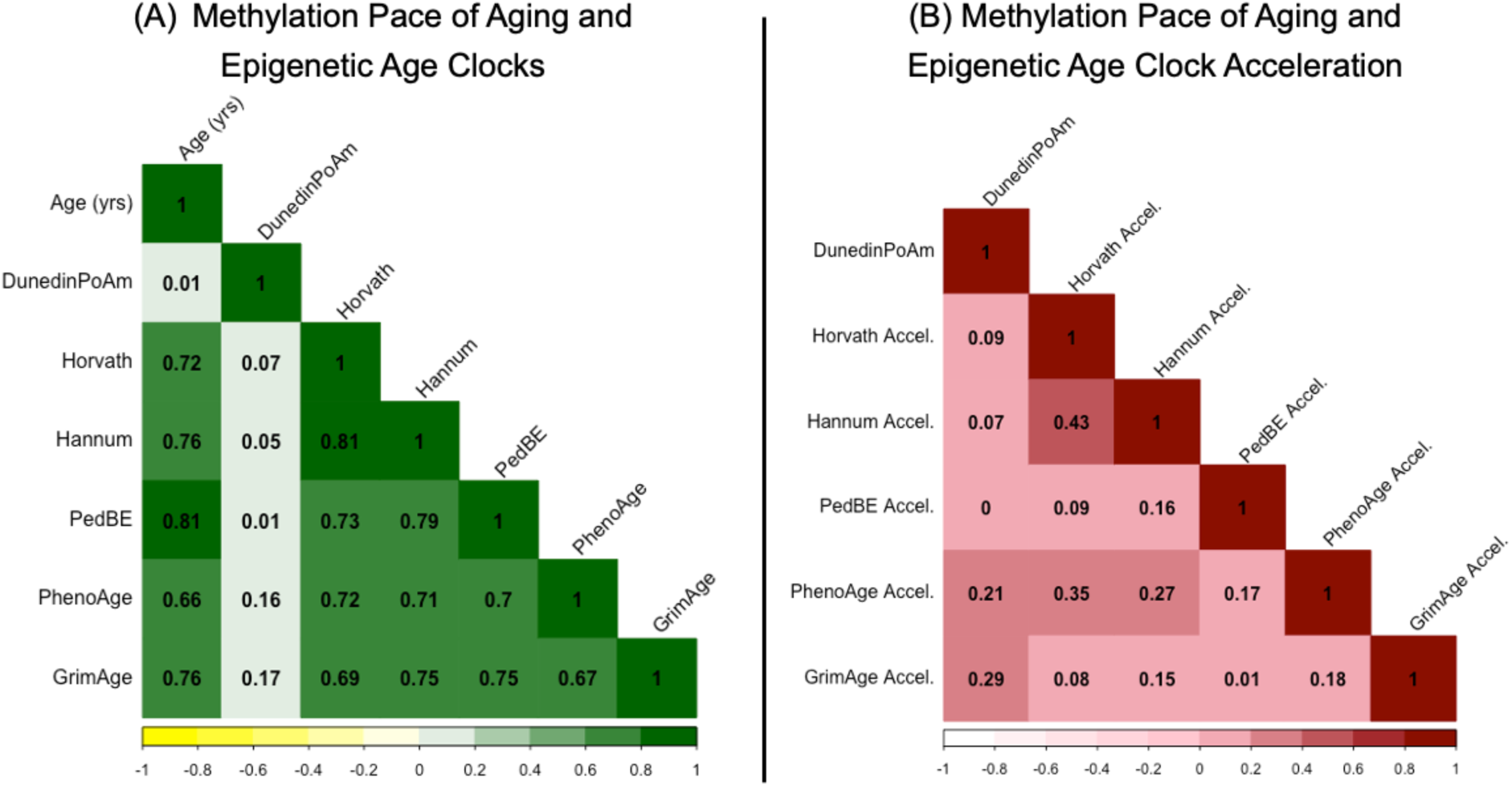
Correlations among DunedinPoAm and five epigenetic clocks. Panel (A) shows the correlation matrix of the methylation Pace of Aging (DunedinPoAm) and epigenetic age clocks with chronological age after residualizing for array, slide, and cell composition. Panel (B) shows the correlation matrix of the methylation Pace of Aging (DunedinPoAm) and epigenetic age acceleration after residualizing for chronological age, array, slide, and cell composition.

For analysis, we regressed clock values on children’s chronological age to compute age-acceleration-residuals. These residuals are interpreted as how much more or less aging has occurred in a person as compared to the expectation based on their chronological age. DunedinPoAm was moderately correlated with the PhenoAge and GrimAge clock residuals (PhenoAge *r* =0.21, Grim-Age *r* = 0.29) and weakly correlated with the Horvath and Hannum clock residuals (Horvath *r* =0.09, Hannum *r* = 0.07). Correlations are reported in panel B of Figure 3.

We repeated analysis of socioeconomic status, replacing DunedinPoAm with each of the epigenetic clock residuals in turn. In contrast to results for DunedinPoAm, epigenetic clocks were not associated with socioeconomic disadvantage and, in two cases, associations were in the opposite direction expected (range of *r’s* = -0.025 – 0.050; Table 1).

**Table 1.** Associations between family- and neighborhood-level socioeconomic disadvantage and six saliva DNA-methylation indices.

|  | Family-level<br>socioeconomic disadvantage |  |  | Neighborhood-level<br>socioeconomic disadvantage |  |  |
| --- | --- | --- | --- | --- | --- | --- |
|  | *r* | 95% CI | *p* | *r* | 95% CI | *p* |
| DunedinPoAm | 0.176 | 0.080 – 0.270 | 0.001 | 0.176 | 0.073 – 0.277 | 0.001 |
| Horvath Accel. | 0.017 | -0.046 – 0.080 | 0.613 | 0.024 | -0.053 – 0.114 | 0.464 |
| Hannum Accel. | -0.025 | -0.080 – 0.036 | 0.427 | 0.034 | -0.037 – 0.110 | 0.290 |
| PedBE Accel. | -0.006 | -0.070 – 0.047 | 0.824 | 0.046 | -0.003 – 0.109 | 0.089 |
| PhenoAge Accel. | 0.014 | -0.052 – 0.072 | 0.688 | 0.030 | -0.054 – 0.120 | 0.394 |
| GrimAge Accel. | 0.032 | -0.023 – 0.088 | 0.277 | 0.050 | -0.011 – 0.125 | 0.106 |

Standardized regression coefficients (*r*) and 95% confidence intervals (CIs) calculated by regressing DNA-methylation measures on family-level socioeconomic disadvantage and neighborhood-level socioeconomic disadvantage, separately.

## Discussion

We analyzed saliva DNA-methylation data from children and adolescents participating in the Texas Twin Project to test associations of childhood socioeconomic disadvantage with a novel DNA-methylation measure of the pace of biological aging, DunedinPoAm. We found that children and adolescents growing up under conditions of higher socioeconomic disadvantage exhibited a faster methylation pace of aging as measured by DunedinPoAm. Faster methylation pace of aging in children with higher body mass index and more advanced pubertal development for their age, but these covariates did not account for observed socioeconomic differences. Our results suggest that DunedinPoAm is useful as a salivary biomarker that not only reflects biological aging in adulthood, as was previously established (24), but is also sensitive in real-time to social determinants of health experienced during childhood.

Our analysis of racial/ethnic group differences found that Latinx-identifying children, who faced substantially higher mean levels of both family-level and neighborhood-level cumulative socioeconomic disadvantage, exhibited faster DunedinPoAm compared to both White and Latinx-White identifying children. These group differences in DunedinPoAm were statistically accounted for by differences in cumulative socioeconomic disadvantage. Thus, our findings are consistent with the observations that racial/ethnic differences in socioeconomic disadvantage are an important contributor to racial/ethnic disparities in health (47), Importantly, racial/ethnic disparities in adult health typically persist, although reduced, across all levels of socioeconomic status, for example due to race-based discrimination (32,48,49).

In contrast to findings for DuedinPoAm, analysis of several published DNA-methylation clocks yielded null associations with socioeconomic disadvantage. These clocks showed the expected associations with children’s chronological age (*r* = 0.66-0.81 in an age range of 8-18 years). However, the difference between participants’ epigenetic ages and their chronological ages, *i.e.*, epigenetic age acceleration, did not differ between children experiencing different levels of socioeconomic risk, consistent with results from a recent meta-analysis (23). Measures of epigenetic age acceleration also appear to be less sensitive to racial/ethnic group differences in childhood and adolescence (22). These results suggest that pace-of-aging measures such as DunedinPoAm may prove more sensitive to health damaging effects of socioeconomic deprivation and of racialized disparities in socioeconomic status, particularly in studies focusing on the early lifecourse.

We acknowledge limitations. First, we measured methylation in saliva DNA (which comes a mixture of buccal cells and blood cells), whereas many of the epigenetic indices we analyze were developed using blood or other tissues. We used DNA-methylation algorithms to make statistical adjustment for the cellular composition of the saliva samples. Increasing confidence in our findings, they replicate results for DunedinPoAm and epigenetic clocks from blood DNA methylation measured in 18-year-olds (24). Second, because this cohort is still young, we do not know if faster aging observed in childhood will translate into higher disease risk later in life. Continued follow-up is needed. Fourth, our study utilized an observational design. It cannot definitively establish DunedinPoAm as the mediating link in a causal process through which adverse childhood exposures translate into aging-related health gradients. Natural experiment studies and analysis of randomized trials testing social programs are needed (50). Finally, the biology that causes variation in DunedinPoAm and the epigenetic clocks remains poorly understood. Epigenetic changes are understood to be core features of the biological process of aging (51,52). Yet, the methylation measurements we studied are only correlates of the unobserved processes of biological aging, not direct observations of it (53).

Within the bounds of these limitations, our findings have implications for theory and future research. Theory and evidence from animal models suggest epigenetic changes are a mediator of early-life adversity’s effects on aging-related health decline (54). However, human studies following-up specific mechanisms identified in animals have yielded equivocal results (55). Our results add to evidence that one mechanism linking early adversity with adult disease might be acceleration of the pace of biological aging.

Future research can take advantage of measurements such as DunedinPoAm to further elucidate how aging processes may be accelerated in at-risk young people and to test if and how such accelerated aging may be modified. Childhood interventions to improve equitable access to healthy food, lower family stress, neighborhood safety, and greenspace have the potential to improve concurrent and lifelong health (6). However, childhood interventions antedate the onset of adult disease by decades. This long gap has motivated interest in biological measures that can serve as surrogate endpoints for assessing the effectiveness of programs and policies that aim to improve lifelong health by promoting positive child development. Our results suggest that salivary DNA-methylation measures of pace of aging may provide a surrogate or intermediate endpoint for understanding the health impacts of such interventions. Such applications may prove particularly useful for evaluating the effectiveness of health-promoting interventions in at-risk groups.

## Acknowledgments

We gratefully acknowledge all participants of the Texas Twin Project. This research was supported by NIH grant R01HD083613 and R01HD092548. LR is supported by the German Research Foundation (DFG). KPH and EMTD are Faculty Research Associates of the Population Research Center at the University of Texas at Austin, which is supported by a grant, 5-R24-HD042849, from the Eunice Kennedy Shriver National Institute of Child Health and Human Development (NICHD). KPH and EMTD are also supported by Jacobs Foundation Research Fellowships. DWB is supported by grants from the Russel Sage Foundation (1810-08987) and the National Institute on Aging (R01AG066887, R01AG061378) and by the Canadian Institute for Advanced Research and the Jacobs Foundation.

## Legends for tables and figures

**Table 1. Associations between socioeconomic disadvantage and epigenetic indices of aging.**

**Figure 1. Associations between family-level and neighborhood-level socioeconomic disadvantage and the methylation Pace of Aging (DunedinPoAm)**. DunedinPoAm and socioeconomic disadvantage values are in standard deviation units. Higher values indicate a methylation profile of faster biological aging. Regression is estimated from linear mixed effects model that accounts for nesting of children within families. The shaded areas represent the smoothed lower and upper 95% confidence intervals.

**Figure 2. Group differences in the methylation Pace of Aging (DunedinPoAm) between children identifying as White only, Latinx only, and both Latinx and White.** DunedinPoAm values are in standard deviation units. Higher values indicate a methylation profile of faster biological aging. Regression is estimated from linear mixed effects model that accounts for nesting of children within families. The boxplot displays group DunedinPoAm differences in the mean (black circle), standard errors of the mean (error bars), and the first and third quartiles (lower and upper hinges). Group differences were significant at the alpha=0.05 threshold without adjustment for differences in socioeconomic disadvantage between groups (left panel), but were no longer significantly different from zero when controlling for family-level and neighborhood-level disadvantage (right panel).

**Figure 3. Correlation between methylation Pace of Aging (DunedinPoAm) and five epigenetic clocks.** Panel (A) shows the correlation matrix of the methylation Pace of Aging (DunedinPoAm) and epigenetic age clocks with chronological age after residualizing for array, slide, and cell composition. Panel (B) shows the correlation matrix of the methylation Pace of Aging (DunedinPoAm) and epigenetic age acceleration after residualizing for chronological age, array, slide, and cell composition.

## Supplemental Information

### DNA methylation preprocessing

DNA methylation preprocessing was conducted with the ‘minfi’ package (56). Prior to calculation of methylation profiles, CpG probes with detection p > 0.05 and fewer than 3 beads in more than 1% of the samples, probes in cross-reactive regions, and those containing a SNP with minor allele frequency above 0.01 within 10 base pairs of the single base extension position or at the CpG interrogation were excluded (57). Methylation values were normalized with noob background correction as implemented by minfi’s “preprocessNoob” (58). 8 participants were excluded for poor performing probes: they showed low intensity probes as indicated by the log of average methylation <11 and their detection p was > 0.05 in >10% of their probes.

DNA methylation differs between cell tissue types, thus it is important to residualize for cell composition. Cell composition was estimated in two ways. First, the blood-based package “FlowSorted.Blood.450k” R package estimates CD8 T cells, CD4T cells, natural killer cells, B cells, monocytes, and granulocytes (59). Second, we estimated 5 cell types using the tissue reference-free method by Houseman et al. (60) as implemented in the “RefFreeCellMixArray” R package. The association of DunedinPoAm residualized with the blood-based control and DunedinPoAm residualized with the reference-free method was very strong (*r* = 0.97, 95% CI = 0.953 – 0.991, *p* < 0.001). We therefore report results of methylation profiles residualizing for cell composition using the blood-based estimation. Methylation profiles were also residualized for array and slide; all samples came from the same batch.

